# Aberrant glycosylation of anti-SARS-CoV-2 IgG is a pro-thrombotic stimulus for platelets

**DOI:** 10.1101/2021.03.26.437014

**Authors:** Alexander P Bye, Willianne Hoepel, Joanne L Mitchell, Sophie Jégouic, Silvia Loureiro, Tanya Sage, Steven de Taeye, Marit van Gils, Neline Kriek, Nichola Cooper, Ian Jones, Jeroen den Dunnen, Jonathan M Gibbins

## Abstract

A subset of patients with COVID-19 become critically ill, suffering from severe respiratory problems and also increased rates of thrombosis. The causes of thrombosis in severely ill COVID-19 patients are still emerging, but the coincidence of critical illness with the timing of the onset of adaptive immunity could implicate an excessive immune response. We hypothesised that platelets might be susceptible to activation by anti-SARS-CoV-2 antibodies and contribute to thrombosis. We found that immune complexes containing recombinant SARS-CoV-2 spike protein and anti-spike IgG enhanced platelet-mediated thrombosis on von Willebrand Factor *in vitro*, but only when the glycosylation state of the Fc domain was modified to correspond with the aberrant glycosylation previously identified in patients with severe COVID-19. Furthermore, we found that activation was dependent on FcγRIIA and we provide *in vitro* evidence that this pathogenic platelet activation can be counteracted by therapeutic small molecules R406 (fostamatinib) and ibrutinib that inhibit tyrosine kinases syk and btk respectively or by the P2Y12 antagonist cangrelor.

## Introduction

COVID-19 is more likely to progress to a severe, life threatening condition in patients with pre-existing cardiovascular disease and is associated with dysregulated haemostasis and a high incidence of venous and arterial thromboembolism [1–3]. Emboli in the pulmonary arteries and microthrombi containing fibrin and platelets in the pulmonary microvasculature of COVID-19 patients have been identified at post-mortem [4] and are thought to contribute toward development of acute respiratory distress syndrome (ARDS). It is now believed that multiple factors contribute to the thromboinflammatory state that results in high rates of thrombotic complications. Evidence has indicated the presence of activated vascular endothelial cells, macrophages, platelets, neutrophils and an activated coagulation system in critically ill COVID-19 patients. The mechanistic trigger that causes the changes that accompany an increase in severity in a subset of patients is still the subject of intense research. However, the disparity between the time of peak viral load at 5-6 days after the onset of symptoms and occurrence of ARDS after 8-9 days, implicate an excessive immune response rather than direct actions of the virus itself [5].

Further evidence that the adaptive immune response is disturbed in severely ill COVID-19 patients has been provided by a study that found high levels of extrafollicular B cell activation in critically ill patients which correlates with increased morbidity, antibody titres and levels of inflammatory biomarkers [6]. Other studies have also noted the strong association of high antibody titres with disease severity and survival [7, 8]. However, antibodies of severely ill COVID-19 patients have qualitative as well as quantitative differences compared to those with mild illness. Anti-spike IgG in serum samples from severely ill COVID-19 patients were found to have low levels of fucosylatation and elevated galactosylation in the Fc domain [9, 10].

Platelets express the antibody receptor FcγRIIA, but it is not known if immune complexes containing afucosylated IgG might activate platelet FcγRIIA. Clustering of platelet FcγRIIA induced by ligand binding triggers intracellular signalling via syk activation and promotes granule secretion and integrin αIIBβ3 activation [11]. Therefore, activation of FcγRIIA by afucosylated anti SARS-CoV-2 spike IgG might further exacerbate thromboinflammation in critically ill COVID-19 patients.

In this study we investigate the effects of low fucosylation and high galactosylation of antispike IgG on platelet activation to find the significance of aberrant IgG glycosylation identified in critically-ill COVID-19 patients on platelet-mediated thrombus formation. We found that potent activation of platelets by immune complexes containing SARS-CoV-2 spike and anti-spike IgG only occurs when the IgG expresses both low fucosylation and high galactosylation in the Fc domain and an additional prothrombotic signal, we used vWF, is also present. Enhanced platelet activation and thrombosis, measure *in vitro*, was sensitive to FcγRIIA inhibition and small molecules inhibitors of syk, btk and P2Y12, suggesting that these therapeutic strategies might reduce platelet-mediated thrombosis in critically-ill COVID-19 patients.

## Materials and Methods

### Spike protein

The sequence of SARS-CoV-2 S1 was obtained from the cloned full-length S sequence and was cloned into the expression vector pTriEx1.1 (EMD Millipore, UK) and characterised as described previously [12]. Sf9 cells were transfected with the baculovirus expression vector FlashBAC Gold (Oxford Expression Technologies, UK) and with the SARS-CoV2-S1 transfer vector to produce recombinant baculovirus. Large-scale protein expression was performed by infecting 1L of T.nao38 cells with a high titre stock of the recombinant baculovirus and incubated for 3-5 days at 27°C. After incubation the supernatant containing the secreted protein was harvested, clarified by centrifugation at 4,300xg for 20min and filtered through a 0.45μm filter. The clear supernatant was supplemented with 0.5nM nickel sulphate before being loaded onto the Bio-Scale Mini Profinity IMAC Cartridge (Bio-Rad, UK). The elution was carried out at a flow rate of 2.5 ml/min with a gradient elution of 0.05-0.25M imidazole over 60 min.

### Recombinant anti SARS-CoV-2 IgG

COVA1-18 IgG was produced in 293F cells as described previously [13]. Antibodies with modified glycosylation states were generated and validated as previously described [14]. Validation of the modifications made to COVA1-18 glycosylation are included in table 1.

**Table 1.**
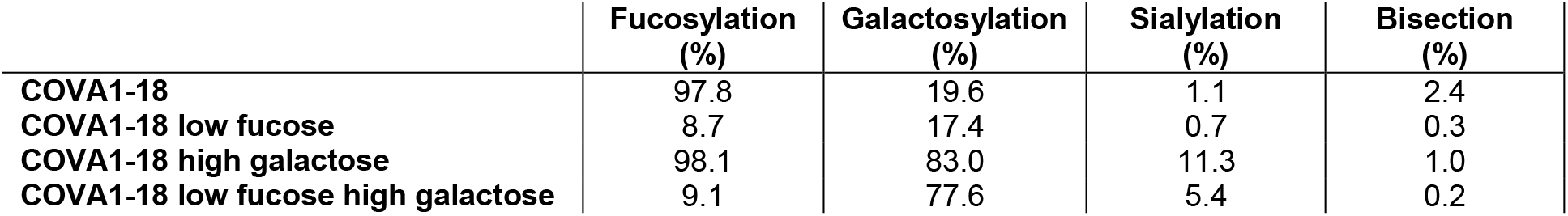
Glycosylation of WT and modified COVA1-18 IgG.

### Blood Preparation

Blood samples were obtained from healthy donors that had given informed consent and using procedures approved by the University of Reading Research Ethics Committee and collected into vacutainers containing 3.8% (w/v) sodium citrate. Platelet-rich plasma (PRP) was prepared by centrifuging whole blood at 100g for 20 minutes. Washed platelets were prepared by adding acid citrate dextrose to PRP and centrifuging at 350g for 20 minutes and resuspending the platelet pellet at 4 × 10^8^ cells/mL in Tyrode’s buffer (NaCl 134 mM, KCl 2.68 mM, CaCl_2_ 1.80 mM, MgCl_2_ 1.05 mM, NaH_2_PO_4_ 417 μM, NaHCO_3_ 11.9 mM, Glucose 5.56 mM, pH 7.4).

### Platelet adhesion assay

Glass 8 well microslides (Ibidi, Munich, Germany) were coated with 5μg/ml recombinant SARS-COV-2 spike protein for 60 minutes at 37°C, washed and then blocked with 10% fetal calf serum (FCS) for 1h at 37°C. The slides were then washed and treated with 10μg/ml COVA1-18 recombinant, monoclonal anti-SARS-COV-2 antibodies (see above) for 1h at 37°C. Washed platelets at 2 × 10^7^ cells/mL were incubated on the slides for 1h at 37°C. Non-adherent platelets were washed off with Tyrode’s buffer and then slides were fixed with 10% formyl saline for 10 minutes. Wells were then washed and the platelets were labelled with 2μM DiOC_6_. Fluorescence images of adherent platelets were captured with the 20× objective lens of a confocal Ti2 microscope (Nikon).

### In vitro thrombus formation

Thrombus formation experiments were performed using microfluidic flow chips (Vena8, CellixLtd, Dublin, Ireland) coated with 5μg/ml recombinant SARS-COV-2 spike protein for 60 minutes at 37°C, washed and then blocked with 10% fetal calf serum (FCS) for 1h at 37°C.

The slides were then washed and treated with 10μg/ml COVA1-18 recombinant, monoclonal anti-SARS-COV-2 antibodies (see above) for 1h at 37°C and then 2U/ml vWF for 1h. Thrombus formation was measured by perfusing citrated whole blood through the flow chambers at 1000s^-1^ for 6 minutes before fixing with 10% formyl saline, staining with 2μM DiOC6 and then imaged by acquiring z-stacks using the 20× objective lens of a confocal Ti2 fluorescence microscope (Nikon).

### Light transmission aggregometry

Aggregation was measured in washed platelets in half-area 96-well plates (Greiner) by shaking at 1200 rpm for 5 minutes at 37°C using a plate shaker (Quantifoil Instruments) after stimulating with collagen at a range of concentrations. Changes in light transmittance of 405 nm was measured using a FlexStation 3 plate reader (Molecular Devices).

### Flow cytometry measurement of fibrinogen binding

Measurements of fibrinogen binding were performed using washed platelets pretreated with immune complexes, created by incubating 5μg/ml recombinant SARS-COV-2 spike protein with 10μg/ml COVA1-18 for 60 minutes at 37°C, and then stimulated with 1 μg/mL CRP, 10μM ADP or 1μM TRAP-6 in the presence of fluorescein isothiocyanate–conjugated polyclonal rabbit anti-fibrinogen antibody (Agilent Technologies LDA UK Limited, Cheadle, United Kingdom), and then incubated for 20 minutes in the dark. Platelets were then fixed by addition of filtered formyl saline (0.2% formaldehyde in 0.15M NaCl) and median fluorescence intensities were measured for 5000 platelets per sample on an Accuri C6 Flow Cytometer (BD Biosciences, Berkshire, United Kingdom).

### Statistical methods

Statistical testing as described in figure legends and the results section were performed using GraphPad Prism Software (GraphPad, La Jolla, CA).

## Results

### Aberrant glycosylation of anti-spike IgG enhances in vitro thrombus formation on vWF

Previous studies have found a correlation between the severity of COVID-19 and the glycosylation state of anti-SARS-CoV-2 IgG, with low levels of fucosylation and high levels of galactosylation present predominantly in patients with critical illness [9, 10, 14]. To investigate whether anti-SARS-CoV-2 antibodies with aberrant glycosylation activate platelets, and assess the importance of low focusylation and high galactosylation, we studied adhesion and spreading on coverslips coated with IgG-Spike immune complexes containing recombinant SARS-CoV-2 spike protein and COVA1-18 antibody. Four subtypes of COVA1-18 with different glycosylation states were used in the study; IgG with normal levels of fucosylation and galactosylation (WT), a low fucose IgG (Low Fuc), a high galactose IgG (High Gal) and IgG with both low fucose and high galactose (Low Fuc High Gal). We compared the number of platelets adhered to the four different immune complexes relative to spike protein alone and found no significant difference (Figure 1Ai-ii). Levels of platelet adhesion to spike were not significantly greater than adhesion to FCS which was used as a negative control. Adhesive platelet ligands collagen and cross-linked collagen related peptide (CRP-XL) were included as a positive controls. These results indicated that the immune complexes, regardless of the glycosylation state of the anti-spike IgG, were poor ligands for platelet adhesion.

**Figure 1.**
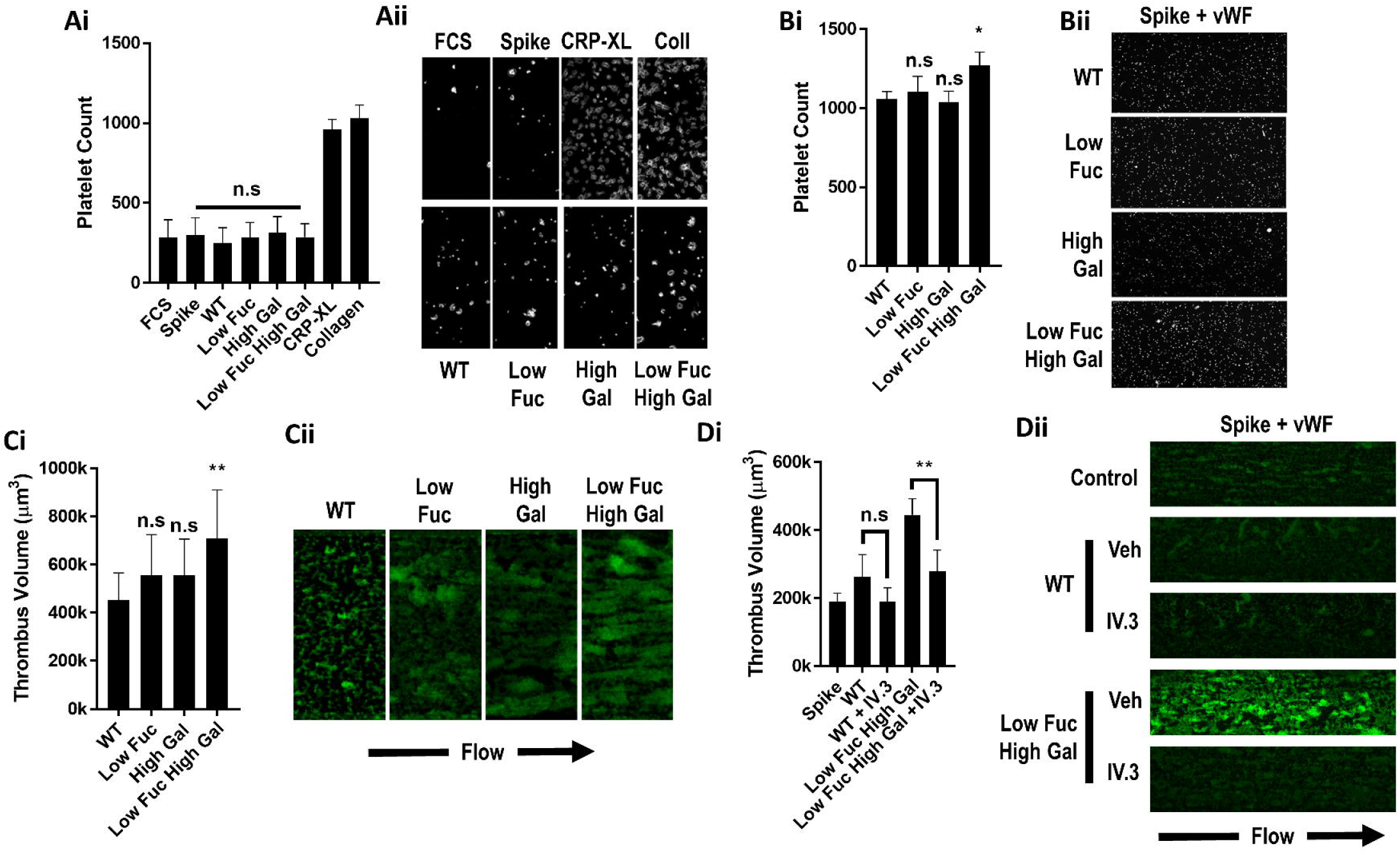
Low fucosylation and high galactosylation of the IgG F tail enhances adhesion to von Willebrand Factor. Platelet adhesion to slides coated with immune complexes containing recombinant SARS-CoV-2 spike protein and COVA1-18 recombinant anti-spike IgG with modified glyocosylation. (Ai) Numbers of platelets adhered to glass slides coated with FCS (negative control), spike protein only or spike protein plus unmodified IgG (WT) or IgG modified to have low fucosylation (low fuc), high galactosylation (high gal) or both (low fuc high gal) and (Aii) representative images of adhered platelets stained with DiOC6. (Bi) Numbers of platelets adhered to vWF plus immune complexes containing spike protein and modified IgGs and (Bii) representative images of adhered platelets. (Ci) Volume of thrombi formed on vWF with immune complexes containing spike and either WT IgG or IgG with modified glyosylation and (Cii) representative images of thrombi stained with DiOC6. (Di) Volume of thrombi formed on vWF plus WT IgG or IgG with low fucosylation and high galactosylation in the presence or absence of 20μg/ml IV.3 and (Dii) representative images of thrombi stained with DiOC6. Values are mean ± s.e.m. Significant differences were tested by 2-way ANOVA with the Tukey multiple comparisons test, * p < 0.05, ** p < 0.01.

Severe COVID-19 is associated with increased levels of many prothrombotic plasma proteins including von Willebrand Factor (vWF), which has been noted as up to five fold higher (to approx. 5U/ml) in COVID-19 patients in intensive care [15–17]. We combined the immune complex coatings with vWF at 2U/ml to simulate a modest increase in plasma vWF levels. The combined immune complex and vWF coating resulted in significantly more platelet adhesion to the Low Fuc High Gal IgG than the WT IgG (Figure 1Bi-ii). Adhesion to IgG with either Low Fuc or High Gal was not significantly different to the WT IgG. This suggested that both hypofucosylation and hypergalactosylation are required for platelet activation by anti-spike IgG immune complexes and that prothrombotic conditions in the circulation of severely ill COVID-19 patients, coupled with dysregulation of IgG glycosylation may result in increased platelet activation.

The increase in platelet adhesion to IgG-spike immune complex and vWF-coated surfaces was modest under static conditions, however, vWF facilitates platelet adhesion to the vascular endothelium in the high shear environment found in arteries and arterioles. To replicate these conditions, we performed thrombus formation experiments in perfusion chambers coated with IgG-spike immune complexes and vWF at a shear rate of 1000s^-1^ (Figure 1Ci-ii). Under these conditions there was no significant difference in the average volume of thrombi formed on vWF combined with Low Fuc or High Gal IgG relative to WT IgG. However, there was a significant increase in thrombus volume formed on vWF in the presence of Low Fuc High Gal IgG immune complexes. This suggests that anti SARS-CoV-2 IgG with aberrant glycosylation of the Fc domain synergises with vWF to enhance thrombus formation. This replicates the synergy observed between platelet receptors that predominantly mediate adhesion, such as GPIb and integrin α2β1, with receptors that strongly activate platelet signalling such as GPVI and CLEC-2 [18].

FcγRIIA is the only Fc receptor expressed in platelets and activates intracellular signalling via Syk activation [19]. We hypothesised FcγRIIA may play the role of a signalling receptor to synergise with the vWF adhesion receptor GPIb to enhance thrombus formation. To test this, whole blood was preincubated with FcγRIIA blocking antibody IV.3 before perfusion through vWF-coated flow chambers with either WT or Low Fuc High Gal immune complexes (Figure 1Di-ii). Blockade of FcγRIIA resulted in a significant reduction in the volume of thrombi formed on the Low Fuc High Gal IgG-containing immune complexes.

### Immune complexes presented in suspension do not potently enhance platelet activation

Platelet aggregability is enhanced in patients with severe COVID19 [20] and we hypothesised that this might be due to the presence of immune complexes containing antispike IgG with aberrant glycosylation of the IgG Fc domain. To test this hypothesis we preincubated recombinant SARS-COV-2 spike protein with the same four COVA1-18 IgG variants used in previous experiments to enable formation of immune complexes in suspension. We then treated washed human platelets with the immune complexes and stimulated with a range of collagen (type I) concentrations to induce aggregation (Figure 2A). We found that none of the immune complexes enhanced the potency (EC50) of collagen-evoked aggregation (Figure 2B) or caused aggregation on their own. We also assessed the ability of the immune complexes to potentiate activation of integrin αIIBβ3 by measuring fibrinogen binding stimulated with ADP, TRAP-6 or CRP-XL by flow cytometry (Figure 2Ci-iii). We observed no significant difference between integrin activation stimulated by agonists in the presence of spike protein alone or immune complexes containing both spike and IgG.

**Figure 2.**
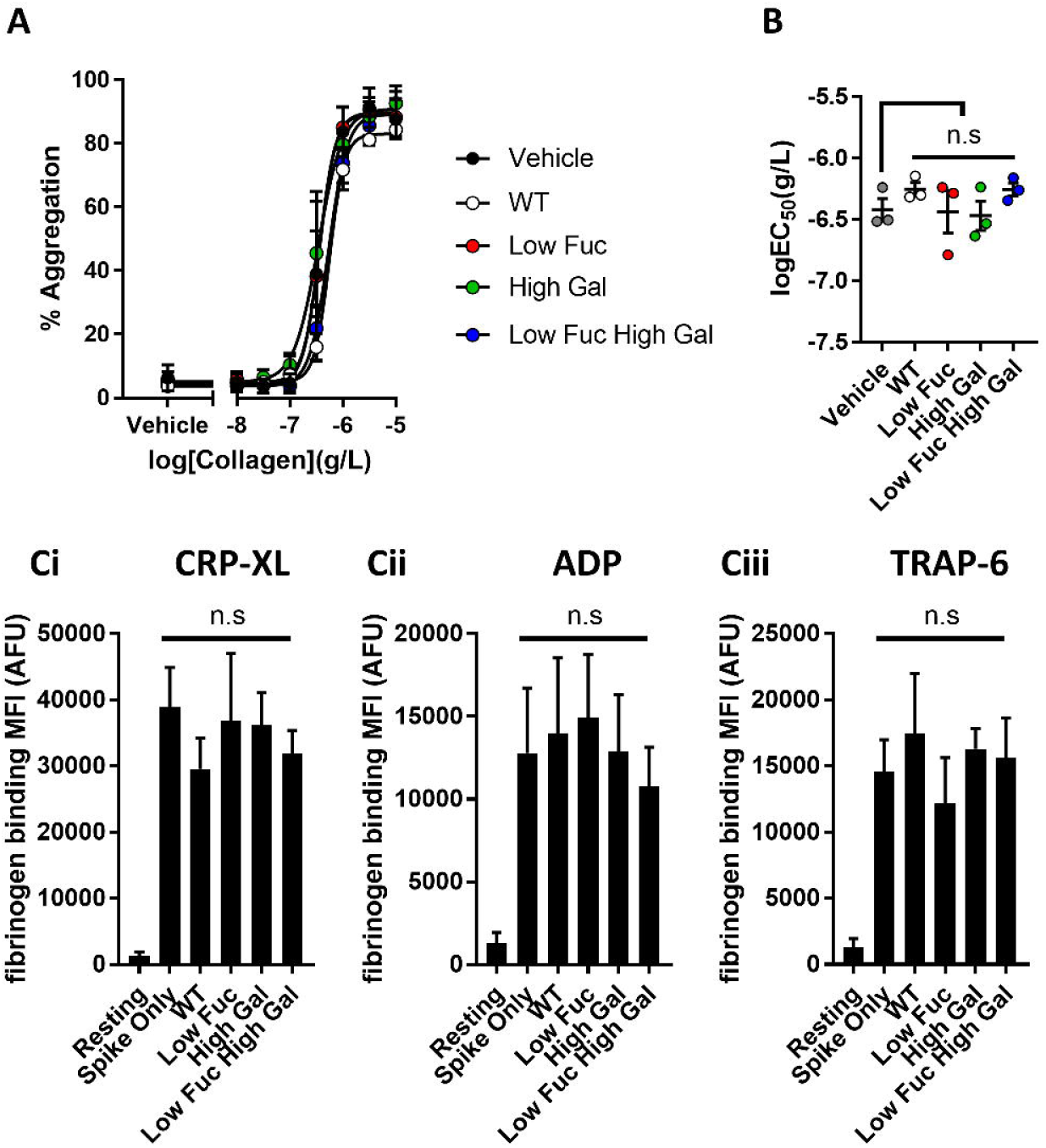
Aggregation and integrin α_IIB_β_3_ activation are unaffected by COVID19 immune complexes. (Ai) Concentration response curves plotting platelet aggregation following stimulation with a range of type I collagen concentrations (from 10μg/ml to 10ng/ml) in the presence of immune complexes containing spike plus WT IgG or IgG with modified glycosylation and (Bi) scatter plots of logEC_50_ for collagen in the presence of the different treatments. Fibrinogen binding to platelets measured by flow cytometry following stimulation with (Ci) 10μM ADP, (Cii) 1μg/ml CRP-XL, (Ciii) 1μM TRAP-6 in the presence of spike only or immune complexes containing WT IgG or IgG with modified gycosylation. Significant differences were tested by 2-way ANOVA with the Tukey multiple comparisons test.

These data suggest that the manner of presentation of immune complexes may be an important part of the mechanism by which they activate platelets. Clustering of FcγRIIA induces intracellular signalling in platelets [19] and it is possible that immune complexes presented in suspension, rather than immobilised on a surface, are below the concentration threshold required to cause clustering of the receptor. Infected endothelial cells may present immobilised immune complexes to platelets in the circulation during severe COVID-19 infection.

### Small molecules drugs targeting syk, btk and P2Y12 inhibit IgG-induced potentiation of thrombus formation on vWF in vitro

To understand the signalling processes underpinning the enhancement of thrombus formation on vWF and Low Fuc High Gal IgG and identify potential treatment strategies to counteract pathogenic platelet activation we studied the effects of small molecule inhibitors (Figure 3). FcγRIIA signals through the tyrosine kinase Syk, so we treated whole blood with the Syk inhibitor R406 which is the active metabolite of the FDA-approved drug fostamatinib. Treatment with R406 significantly reduced the volume of thrombi formed on Low Fuc High Gal IgG (373k ± 42k μm^3^) relative to vehicle (820k ± 172k μm^3^), indicating that activation of Syk is important to the prothrombotic effects of aberrantly glycosylated anti-SARs-CoV-2 IgG and that treatment with fostamatinib might be beneficial for patients with severe COVID19 through suppression of IgG-driven platelet activation. FcγRIIA signals via PLCγ2 [21] and we hypothesised that, similar to that stimulated by the platelet GPVI receptor [22–24], the pathway involved might be dependent on Btk. We therefore treated platelets with the Btk inhibitor ibrutinib, which is an FDA and EMA approved drug for treatment of B cell cancers. We found that ibrutinib treatment reduced the volume of thrombi formed on the Low Fuc High Gal IgG (348k ± 68k μm^3^) to levels similar to the WT IgG (478k ± 76k μm^3^). Finally, platelet activation via FcγRIIA is dependent on ADP secreted dense granules to activate the P2Y12 receptor and provide the positive feedback signalling required for integrin αIIBβ3 activation and aggregation [25]. Blood samples treated with the P2Y12 antagonist cangrelor, were found to reduce the volume of the thrombi formed on Low Fuc High Gal IgG (389k ± 40k μm^3^) to comparable levels to those observed with WT IgG.

**Figure 3.**
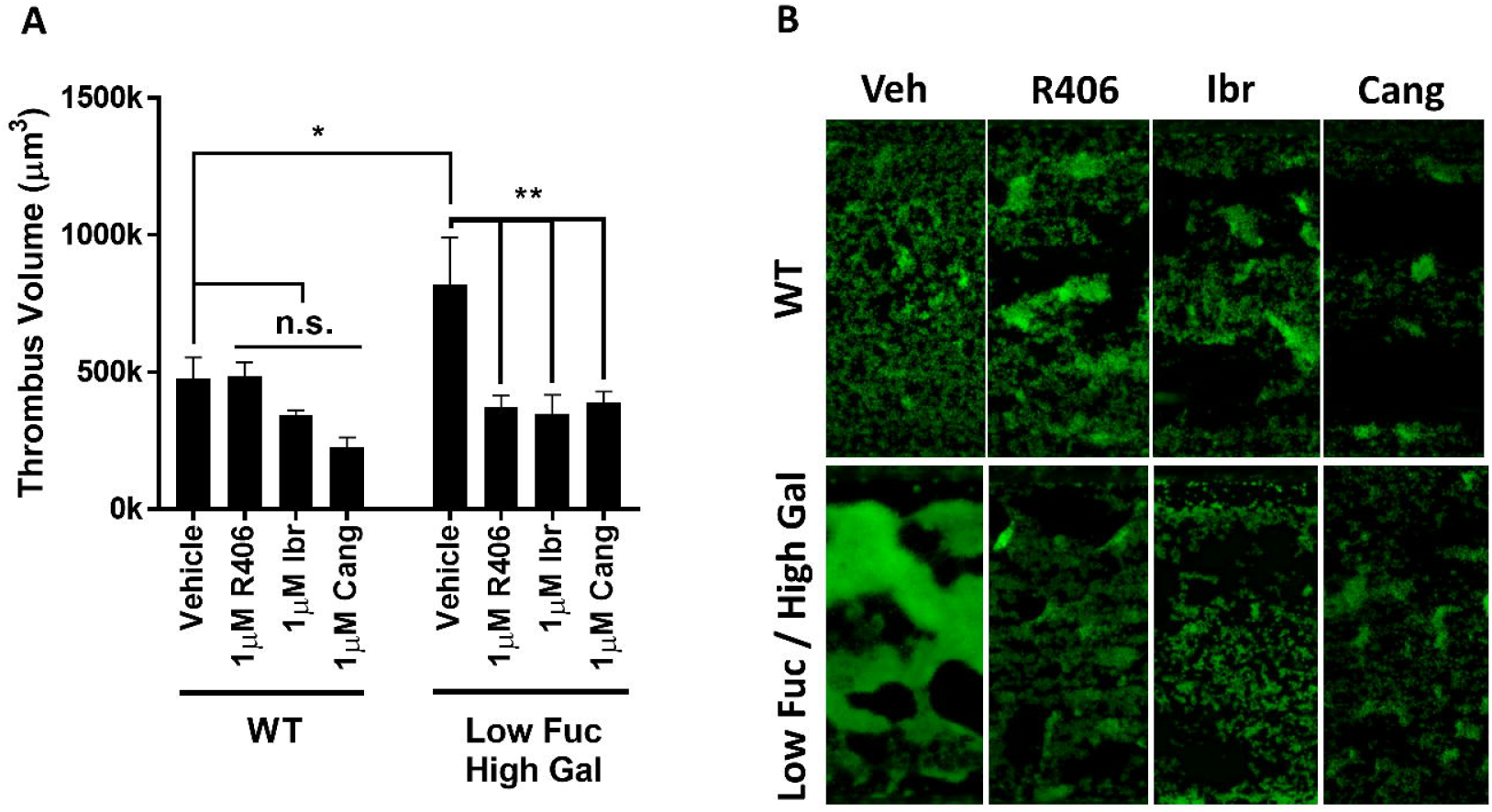
Prothrombotic activity of low fucose, high galactose IgG1 immune complexes is inhibited by Syk, Btk or P2Y12 inhibition. (A) Volume of thrombi formed in perfusion chambers on vWF plus immune complexes containing spike protein plus either WT IgG or IgG modified to have low fucosylation and high galactosylation following treatment with vehicle (DMSO), 1μM R406, 1μm ibrutinib or 1μM cangrelor and (B) representative images of thrombi stained with DiOC6. Values are mean ± s.e.m. Significant differences were tested by 2-way ANOVA with the Tukey multiple comparisons test, * p < 0.05, ** p < 0.01.

## Discussion

There is a growing body of evidence that multiple complications arise in severely ill COVID-19 patients that increase rates of thrombosis. These include damage to vascular endothelial cells following direct infection with SARS-CoV-2, resulting in disruption of barrier function, exposure of subendothelial collagen as well as release of prothrombotic plasma proteins including vWF from activated endothelial cells [16]. The prothrombotic environment is exacerbated by a cytokine storm that may be driven by activation of macrophages by immune complexes containing afucosylated anti-spike IgG [9]. Hypofucosylated, hypergalactosylated IgG has been identified in the plasma of severely ill COVID-19 patients relative to patients with mild COVID-19 infection and correlated with disease severity [9, 14]. In the present study we showed that recombinant anti-spike IgG with low levels of fucosylation and high levels of galactosylation activate platelets to enhance thrombus formation on vWF, which is also elevated in severely ill COVID-19 patients [15–17].

We hypothesise that immune complexes present in the pulmonary circulation of severely ill patients might activate platelets and contribute to high rates of thrombosis (Figure 4). A similar mechanism has been identified in severe H1N1 infection, whereby immune complexes present in the lungs activate platelets via FcγRIIA [26]. The role of aberrant IgG glycosylation in stimulating this response was not investigated, but it has been suggested that afucosylated IgG may be common to immune responses against all enveloped viruses [9].

**Figure 4.**
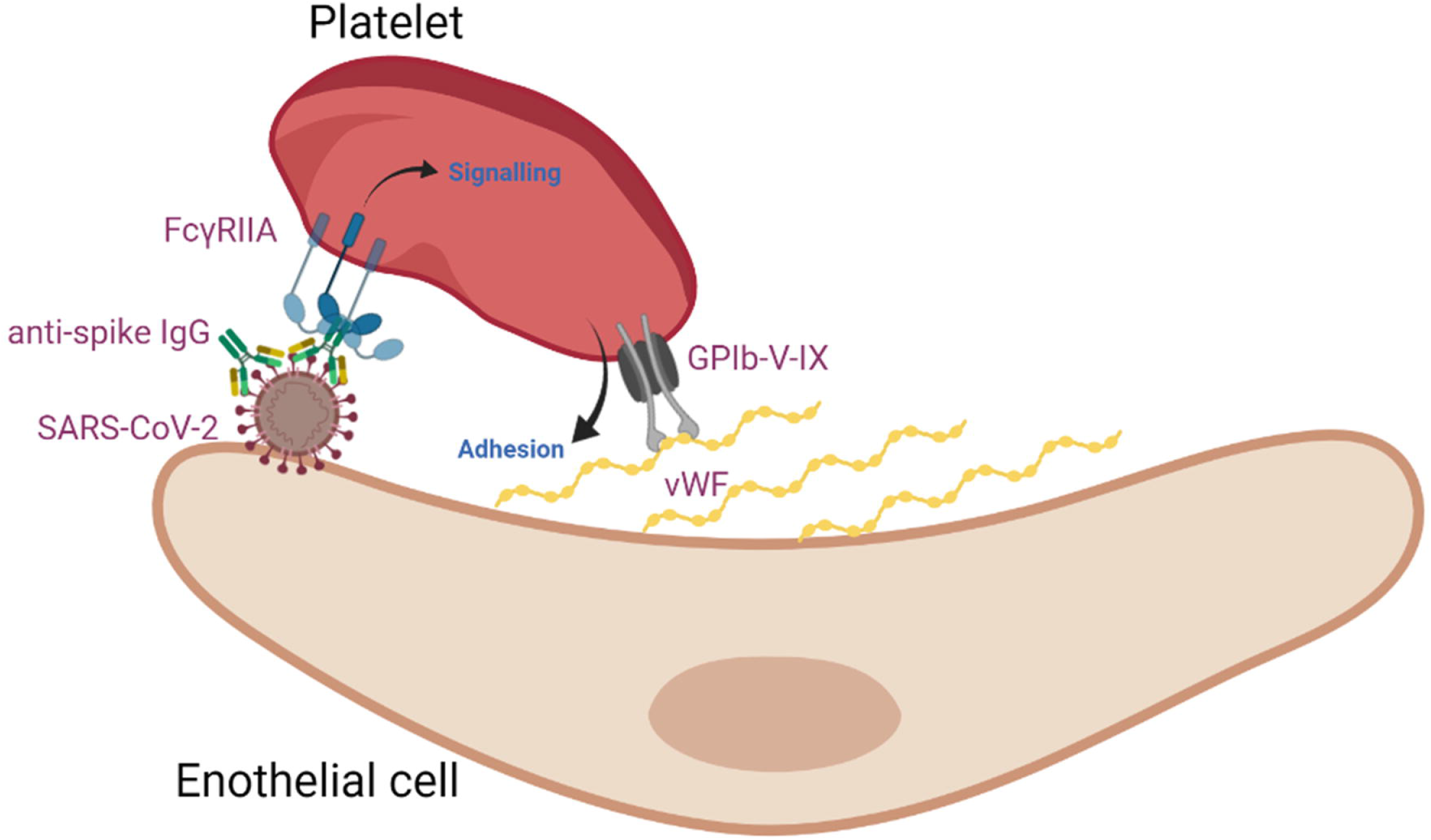
Aberrant glycosylation of anti-spike IgG in immune complexes act in concert with VWF to enhance platelet thrombus formation. SARS-CoV-2 infects vascular endothelial cells, and combined with other inflammatory signals, results in endothelial activation and release of pro-thrombotic factors including VWF. After the onset of adaptive immunity, anti-spike IgG accumulates in the circulation and binds to SARS-CoV-2. In critically ill COVID-19 patients, anti-spike IgG has abnormally low levels of fucosylation and high levels of galactosylation. Immune complexes containing this aberrant glycosylation pattern activate platelet FcγRIIA which stimulate intracellular signals that synergise with the adhesive ligand vWF to promote platelet activation and thrombus formation. Schematic was created with BioRender.com.

The role of platelets in COVID-19 is still emerging but platelet rich thrombi have been identified in both large arteries and microthrombi [4]. Platelets contain many inflammatory mediators within granules that might contribute toward the flood of cytokines present in critically ill COVID-19 patients. There is still scant information regarding the efficacy of antiplatelet drugs in COVID-19 patients but one study has suggested that patients receiving antiplatelet therapy with aspirin prior to hospital admission for COVID-19 appear to be partially protected and have better outcomes, while a separate study found that in-hospital treatment of patients with aspirin reduced mortality [27, 28]. Other non-antiplatelet drugs in trials for COVID-19 therapy have well documented antiplatelet effects. The Bruton’s tyrosine kinase (Btk) inhibitor acalabrutinib has been evaluated in clinical trials on the basis of its potential to block macrophage activation [29], however, the potential of Btk inhibitors to reduce the contribution of platelets to thrombosis in COVID-19 infection have also been noted [30]. We found that the btk inhibitor ibrutinib reversed the enhancement of thrombus formation on vWF caused by IgG with low fucosylation and high galactosylation, supporting the hypothesis that this strategy might have dual benefits on macrophage and platelet activation. The syk inhibitor fostamatinib was identified as a potential COVID-19 therapeutic in a high content screen of drugs that might protect against acute lung injury [31]. The active metabolite of fostamatinib, R406 inhibits release of neutrophil extracellular NETS [32] and macrophage activation induced by plasma from COVID-19 patients [14]. R406 is also known to have potent antiplatelet effects downstream of the GPVI and CLEC-2 receptors [33] and is currently in clinical trials for COVID-19 therapy in the US (NCT04579393) and UK (NCT04581954). We found that R406 reversed the potentiation of thrombus formation on vWF. This suggests that potential COVID-19 therapies such as fostamatinib or acalabrutinib, targeting syk or btk respectively, may be effective not only in limiting the inflammatory response, but also in reducing platelet-mediated thrombosis.

## Acknowledgements

This work was supported by Imperial College NIHR bioresources and grants from the British Heart Foundation (RG/15/2/31224 and RG/20/7/34866), ZonMw (10430 01 201 0008) and an Amsterdam Infection & Immunity COVID-19 grant (24184).

